# Closely related budding yeast species respond to different ecological signals for spore activation

**DOI:** 10.1101/2020.08.11.246728

**Authors:** Samuel Plante, Christian R Landry

## Abstract

Spore activation is one of the most important developmental decisions in fungi as it initiates the transition from dormant and stress resistant cells to vegetative cells. Because in many species mating follows spore activation and germination, signals that trigger this developmental transition can also contribute to species reproductive barriers. Here we examine the biochemical signals triggering spore activation in a natural species complex of budding yeast, *Saccharomyces paradoxus* (lineages *SpA, SpB, SpC* and *SpC*\*). We first demonstrate that we can quantitatively monitor spore activation in these closely related lineages. Second, we dissect the composition of culture media to identify components necessary and/or sufficient to activate spores in the four lineages. We show that, contrary to expectation, glucose is necessary but not sufficient to trigger spore activation. We also show that two of the North American lineages (*SpC* and *SpC*\*) diverge from the other North American (*SpB*) and European (*SpA*) lineages in terms of germination signal as their spore activation requires inorganic phosphate. Our results show that the way budding yeast interpret environmental conditions during spore activation diverged among closely related and incipient species, which means that it may play a role in their ecological differentiation and reproductive isolation.

## Introduction

As part of their life-cycles, fungi produce spores that are specialized for dispersal and survival in harsh environments. In spite of the large diversity of sporulation processes, including the production mode through meiosis or mitosis, fungal spores share important common features. These features include for instance a thick spore wall, the accumulation of protective molecules such as the non-reductive sugar trehalose, and a low metabolic activity (Neiman, 2005). These features increase spore survival-compared to vegetative cells-through long periods of time and difficult environmental conditions, such as extreme temperatures, desiccation and exposure to digestive enzymes (Coluccio et al., 2008; Neiman, 2011). In addition, since fungal spores count for a large proportion of airborne coarse particles (Fröhlich-Nowoisky et al., 2009), most of the fungal-host interactions are mediated by spores (Huang & Hull, 2017; Rieux et al., 2014; Walsh et al., 2019). Notably, *Cryptococcus neoformans* spore properties and specific interaction with macrophages contribute to its virulence after primary contact in alveoli (Botts & Hull, 2010).

Because spores are often associated with harsh environmental conditions, the spore-to-vegetative cell developmental switch and decision is particularly important. Resumption of growth in spores is a multi-step program termed germination, which involves both unique processes (Allen, 1965; Plante & Labbé, 2019; van Leeuwen et al., 2013), and others that are shared with similar transitions such as exit from stationary phase in yeast and G_0_-to-G_1_ transition in mammalian cells. Germination is initiated when spores recognize adequate environmental conditions. Early events of germination that follow its induction are termed exit from dormancy, or spore activation, which we will use interchangeably. These events involve loss of the protective features and gradual gain of vegetative cell characteristics. Since initiation of germination is irreversible (Herman & Rine, 1997; Van Laere et al., 1983), spore activation represents a commitment. Therefore, precise signals are needed for initiation of germination in order to avoid exit from dormancy in conditions incompatible with the growth of vegetative cells.

Fungal spores bear various shapes and ornaments linked to their life-style. For instance, some have structures that ease dispersal by air and insect vectors, and others for adhesion on specific substrates (Calhim et al., 2018; Halbwachs et al., 2015; Pringle et al., 2015). However, we have a poor knowledge of the diversity in nutritional determinants for germination, which are also expected to be ecological-niche specific given the strong selective pressures acting on this decision. Early studies on spore germination using laboratory strains of species such as *Saccharomyces cerevisiæ, Schizosaccharomyces pombe* or *Aspergillus nidulans* identified sugars as an essential and sufficient requirement for activation (d’Enfert, 1997; Herman & Rine, 1997; Shimoda, 1980). Yet, there are some examples of fungal spore germination stimulated by specific compounds or physicochemical properties of their natural niche. Amongst them, *Neurospora crassa* spores are known to germinate following a heat-shock, which has been linked to their propancy to colonize tree remains quickly after a fire (Schmit & Brody, 1975). Root exudate, such as flavonoids, were found to initiate germination of soil-borne fungal spores (Bagga & Straney, 2000), while *Colletotrichum* species and rust-fungi spore germination is promoted by cuticular waxes or plant volatile compounds (French, 1992; Podila et al., 1993). These are examples of germination regulation through specific requirements preventing commitment to germination in other environments than the one these species usually thrive and propagate in.

In species such as *Saccharomyces sp*., mating and return to diploidy usually follow haploid spore germination. Although most matings occur among spores from the same ascii (Tsai et al., 2008), these species do outcross. The possibility of a given cell to outcross with unrelated ones in a given environment will therefore depend on these cells responding to the same spore activation signals. Species of the genus *Saccharomyces* show very little prezygotic isolation (Greig, 2009) but spore activation dynamics has been shown to play a potential role in preventing mating among species (Maclean & Greig, 2008; Murphy & Zeyl, 2012). Some of these species have overlapping geographical ranges and show limited gene exchange despite the absence of prezygotic isolation mechanisms. This is the case for the North American *Saccharomyces paradoxus* species complex. For instance, the geographic distributions of *S. paradoxus SpB* and *SpC*, which diverged about 200,000 years ago, overlap partially. Within their overlapping range is also found a hybrid species *SpC*\*, which most likely originated about 10,000 years ago after the glacier retreated and from admixture between *SpB* and *SpC* (Leducq et al., 2016). Another lineage, *SpA*, originally from Europe but recently introduced in North America, has a geographic range that overlaps with that of *SpB*. All of these lineages can be crossed in the laboratory using haploid strains with selectable markers (Charron, Leducq, & Landry, 2014; Leducq et al., 2016). Laboratory F1 hybrids however show various levels of sterility, which implies postzygotic reproductive isolation (Charron, Leducq, Bertin, et al., 2014; Hénault et al., 2017; Leducq et al., 2016). However, F1 hybrids have not been isolated in the wild, suggesting that prezygotic mechanisms of isolation may exist. Because these strains can be easily mated in the laboratory, if prezygotic isolation mechanisms do exist, they may be linked to ecological cues that trigger spore activation rather than from cell-cell interactions. We know relatively little about the molecules that activate spores in *S. paradoxus*, and even less so if these signals differ among the different lineages. Here, we dissect the spore activation conditions of natural isolates of *S. paradoxus* from these different lineages.

## Material and methods

### Yeast strain

All strains used in this study are listed in table 1. They are natural and non-genetically modified isolates from various locations in north-eastern North America. Lineage was attributed based on whole-genome sequencing (Eberlein et al., 2019; Leducq et al., 2016).

**Table 1 -.** Yeast strains used in this study

| **Systematic name** | **lineage** | **Reference** |
| --- | --- | --- |
| LL2012_001 | *SpA* | Leducq et al. 2016 |
| LL2012_026 | *SpA* | Leducq et al. 2016 |
| LL2014_111 | *SpA* | Leducq et al. 2016 |
| LL2016_119 | *SpA* | Eberlein et al. 2019 |
| LL2016_121 | *SpA* | Eberlein et al. 2019 |
| LL2016_134 | *SpA* | Eberlein et al. 2019 |
| LL2012_009 | *SpB* | Leducq et al. 2016 |
| LL2012_014 | *SpB* | Leducq et al. 2016 |
| LL2012_029 | *SpB* | Leducq et al. 2016 |
| LL2014_076 | *SpB* | Eberlein et al. 2019 |
| LL2014_065 | *SpB* | Eberlein et al. 2019 |
| LL2014_080 | *SpB* | Eberlein et al. 2019 |
| LL2016_022 | *SpB* | Eberlein et al. 2019 |
| LL2016_053 | *SpB* | Eberlein et al. 2019 |
| LL2011_002 | *SpC* | Leducq et al. 2016 |
| LL2011_009 | *SpC* | Leducq et al. 2016 |
| LL2013_138 | *SpC* | Leducq et al. 2016 |
| LL2013_210 | *SpC* | Leducq et al. 2016 |
| LL2011_007 | *SpC* | Leducq et al. 2016 |
| LL2011_011 | *SpC* | Leducq et al. 2016 |
| LL2012_010 | *SpC* | Leducq et al. 2016 |
| LL2012_012 | *SpC* | Leducq et al. 2016 |
| LL2011_005 | *SpC\** | Leducq et al. 2016 |
| LL2011_006 | *SpC\** | Leducq et al. 2016 |
| LL2014_059 | *SpC\** | Eberlein et al. 2019 |
| LL2012_016 | SpC* | Leducq et al. 2016 |
| LL2012_018 | SpC* | Leducq et al. 2016 |
| LL2014_099 | SpC* | Eberlein et al. 2019 |

### Sporulation and spore purification

Sporulation was performed as previously described (Tong & Boone, 2006). Briefly, diploid strains were grown to mid-logarithmic phase in liquid YPD medium (1% yeast extract, 2% tryptone, 2% dextrose). Cells were washed in water and incubated on sporulation medium plates (1% potassium acetate, 0.1% yeast extract, 0.05% dextrose, 2% agar, and 0.01% drop-out mix containing uracil, histidine, leucine and lysine at a mass ratio of 1: 1: 5: 1) for at least 3 days at 25°C. Sporulation was monitored by light microscopy. When sporulation was complete (over 90% ascii observed), cells were digested in water in presence of 0.1% *β*-glucuronidase (Sigma, Darmstadt, Germany) and 10 U zymolyase (Bioshop, Burlington, Canada) at 30°C for at least 3 hours. When ascii were broken, and the remaining vegetatives cells were dead, spores were washed in water before being purified on a Percoll gradient as previously described (Plante & Labbé, 2019). Spores were layered on top of a 50-60-70-80% Percoll gradient (From top to bottom, each layer containing 0.5% Triton X-100). After centrifugation at 7,100g for 20 minutes, the pure spore pellets were washed with water. Spore preparations were stored at 4°C in 0.5% Triton X-100 for a maximum of one week.

### Germination induction

Prior to germination induction, media and spore preparations were separately warmed at 30°C for 10 minutes. Spores were inoculated in germination media at concentration of 2 x 10^6^ spores per ml (OD_595_ around 1.0) and incubated at 30°C. Initial time-point of germination refers to the moment right after inoculation of spores. Minimal synthetic medium used for germination contained 5g/L ammonium sulfate, 0.17% yeast nitrogen base (YNB without ammonium sulfate, without amino acids, Bioshop, Burlington, Canada) and 2% dextrose. Minimal medium components were mixed in different combinaisons using these concentrations. YNB components were used separately at the concentration specified in 1x working solution by the YNB supplier (Bioshop, Burlington, Canada). When indicated, potassium phosphate monobasic in YNB was substituted with 7 mM K_2_HPO_4_ or 7 mM acetate buffer pH 4.8. Optical density (OD_595_) of spore cultures was measured at 10 minutes interval over 6 hours in a plate reader (Tecan Infinite M Nano) which was set to 30°C without agitation. Steepest OD_595_ decrease (ΔOD_min_) was defined as the minimal value obtained when subtracting OD_595_ value of a given time-point to the value of the following time-point along the 6 hours-long measurement.

### Live cell imaging of germinating spores

Spores were seeded in eight-well glass-bottom chamber slides (Sarstedt) coated with 0.05 mg/ml concanavalin A (Millipore Sigma) containing 500 μl YPD. Cell imaging was performed on an Apotome Observer Z1 microscope (Zeiss) equipped with LD PlnN 40x/0.6 objective (Zeiss) and a AxioCam MRm camera (Zeiss). Phase contrast images of spores through the germination process were recorded every 10 minutes over 6 hours at 25°C using the AxioVision software (Zeiss).

### Heat resistance measurement

At the indicated time after germination induction cells were sampled. Half of the cells were inoculated in YPD medium, and the other half was treated at 52°C for 10 minutes in a thermocycler (Eppendorf Mastercycler ProS) before being seeded in YPD. OD_595_ of both treated and untreated cells were measured in a plate reader (Tecan infinite M nano) set at 30°C without shaking for 24h. Area under the curve (AUC) was calculated using the Growthcurver package in R (Sprouffske & Wagner, 2016). Heat resistance value was defined as the ratio of AUC of treated growth curve to AUC of untreated growth curve both obtained over the time required for untreated spore ODs to reach stationary phase.

### Flow cytometry analysis

At the indicated time after germination induction, samples of spores were washed in 30 mM EDTA and diluted to 500 cells per μl in YPD before analysis on Guava easyCyte HT cytometer (EMD Millipore). We recorded FCS and fluorescence signals in the orange emission spectrum (620/52 nm) after excitation with a violet laser (405 nm) of 5000 events.

## Results and Discussion

We examined spore activation in four lineages within the *S. paradoxus* system complex. Given the striking morphological changes accompanying yeast germination (Kono et al., 2005; Sando et al., 1980), we first focused on microscopic analysis of wild spores from a subset of strains. Phase contrast microscopy was used to monitor the morphology of spores from strains LL12_001 (*SpA* lineage), LL12_009 (*SpB* lineage), LL11_011 (*SpC* lineage) and LL11_006 (*SpC*\* lineage) after induction of germination in standard growth medium known to trigger spore activation in *S. cerevisiae* (YPD) (Figure 1A). Initially, resting spores were spherical cells, and 160 to 180 minutes after induction, round spores appeared to have increased in size, before their shape elongated. At 300 to 340 minutes post-induction, the first budding events were observed. Bud emergence is a hallmark for germination completion in *Saccharomyces* species (Joseph-Strauss et al., 2007; Kloimwieder & Winston, 2011; Stelkens et al., 2016). Overall, germination of wild *S. paradoxus* ascospores seems to be very similar morphologically and timely between lineages and to what was previously reported in *S. cerevisiae* (Kono et al., 2005).

**Figure 1 -.**
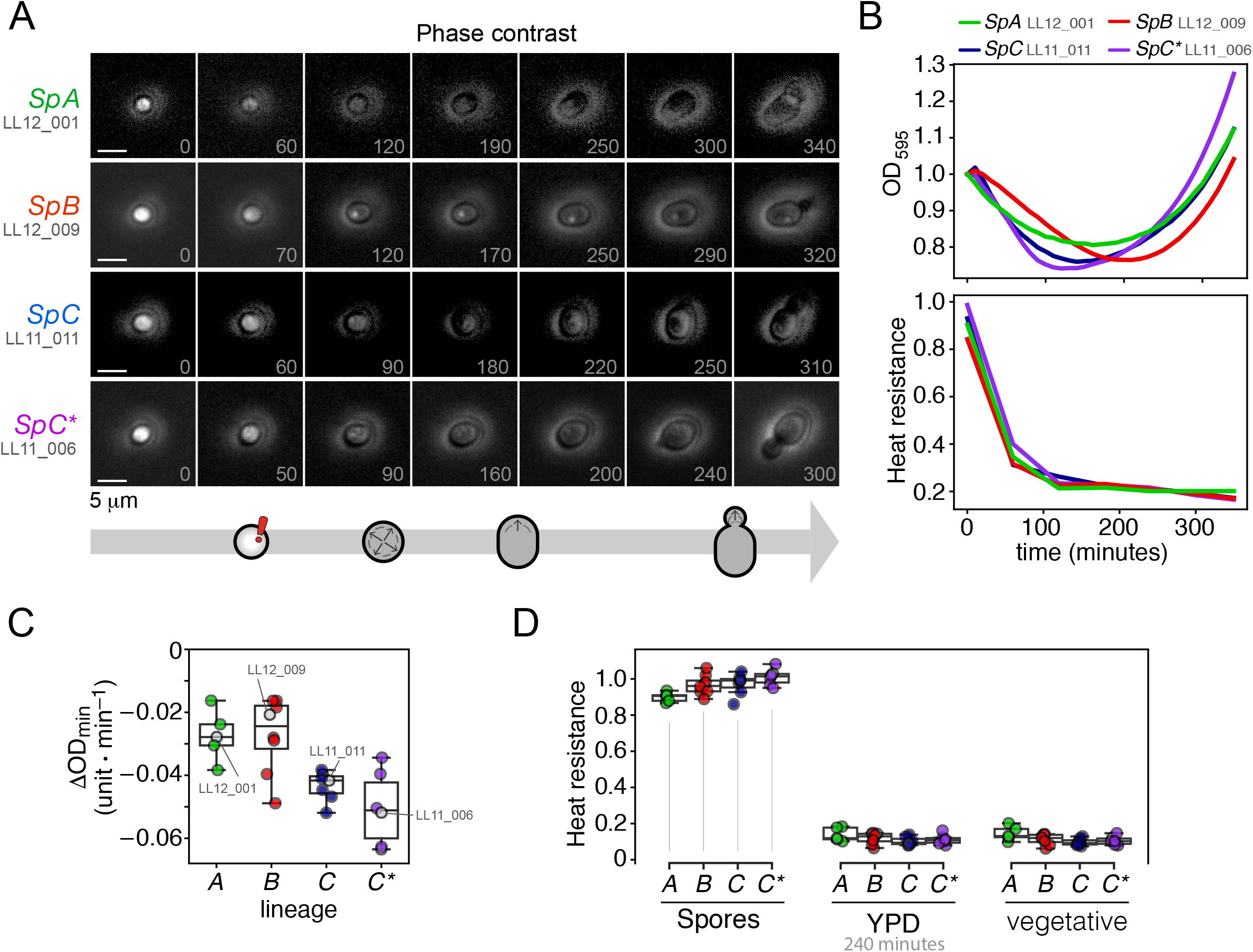
Measurements of spore activation in *Saccharomyces paradoxus*. A) Phase contrast microscopy images of a *SpA*, *SpB*, *SpC* and *SpC*\* spores at the indicated time (in minutes) after germination induction in YPD. Each column depicts precise morphological stages as defined by (Herman & Rine, 1997). Key stages are illustrated at the bottom; the exclamation mark indicates spore activation, arrows within spore indicate growth stage (swelling, elongation and budding respectively). Scale bars represent 5μm. B) *SpA* (green), *SpB* (red), *SpC* (blue) and *SpC\** (purple) spore germination was induced in YPD and activation was measured.Shown are representative data of 4 replicates. The top panel shows OD_595_ at 10 minutes intervals until 360 minutes post-induction. The bottom panel shows resistance to 52°C treatment for 10 minutes at 0, 60, 120, 180, 240, 300 and 360 minutes post-induction (see methods). C) and D) show extension to a larger number of strains of each lineage. C) Steepest OD_595_ decrease (in units per minute) recorded for 28 strains (6 *SpA*, 8 *SpB*, 8 *SpC* and 6 *SpC*)* after germination induction in YPD. D) Heat resistance for spores from 28 strains at initial 0h time-point (spores), at 240 minutes after germination induction in YPD (YPD), or of vegetative cells at mid-log phase (vegetative).

Morphological changes appeared relatively synchronously in the population so we wanted to quantify the synchronicity of spore activation. To do so, we analysed the same *SpA*, *SpB*, *SpC*, and *SpC*\* spores through germination in YPD by flow cytometry. Plotting the forward scatter (FSC) *vs* fluorescent signal of the spore population effectively allows us to differentiate between stages of germination through time (Figure S1). At the initial time point, dormant spores cluster tightly at low FCS and low fluorescent signals. The spore population synchronously gained fluorescence within 2h post-induction, then gained FCS signal until 6h time-point. Since cytometric analysis was very similar in all four strains (Kruskal-Wallis test, significant only for fluorescence at 0h *p* = 0.011), we were confident that our procedure allows similar synchronous induction of germination.

We noted that resting spores appear bright under phase contrast microscopy (figure 1A). Highly refractile spores quickly darkened after germination induction (figure 1a). Bright-phase to dark-phase transition is associated with the exit from dormancy, known as spore activation (germination *per se*). As previously reported in *S. cerevisiæ* and *S. pombe* (Hatanaka & Shimoda, 2001; Rousseau et al., 1972), loss of refractility in individual spores happened in parallel to a decrease in optical density at 595 nm (OD_595_) of spore culture (Figure 1B). Minimal OD_595_ value was reached at 100-to 200-minutes time-point before further outgrowth of spores caused increase of OD_595_. We extended the quantification of germination of spores to 28 wild *S. paradoxus* strains (6 *SpA*, 8 *SpB*, 8 *SpC*, and 6 *SpC*\*). We measured the steepest diminution in OD_595_ (termed ΔOD_min_, Figure 1C) for each strain. Since ΔOD_min_ correlated well with minimal OD_595_ reached (Figure S2, r^2^ = 0.91, *p* = 1×10^-10^), we focused on ΔOD_min_ to describe the OD curve shape.

Drastic alteration of spore surface and content that occur at the onset of spore germination results in increase of its sensibility to many environmental stresses, including heat shock (Rousseau et al., 1972; Smits et al., 2001). This hallmark was previously used as a marker for yeast germination (Joseph-Strauss et al., 2007). We observed that *S. paradoxus* spores from all lineages were 8 to 10-fold more resistant than their respective exponentially growing vegetative cells (figure 1d, *p* < 1×10^-5^ Kruskal-Wallis test). Resistance of spores decreased rapidly after their inoculation in YPD. Within 120 minutes, spore heat resistance dropped to 23 to 28% of initial resistance, and at 240 minutes it stayed as low as 21 to 25% of initial resistance. Heat resistance of all ascospores at the 240-minute time-points was statistically similar to that of their respective vegetative cells (Figure 1d, paired t-test). This timing is consistent with the timing of initial budding observed by microscopy. Spore culture ΔOD_min_ and heat resistance decrease therefore both adequately track the ascospore activation of wild *S. paradoxus*.

Using these measurements of spore activation, we undertook investigation of the nutritional determinants of germination. Studies on many other fungi report that a fermentable carbon source (dextrose being preferred) is a major germination cue (Hayer et al., 2013; Osherov, 2009; Shimoda, 1980). Therefore, we first tested spore activation in YP medium in which dextrose was omitted. In this medium, ΔOD_min_ values were higher, meaning that there was almost no OD_595_ decrease in YP compared to YPD (containing dextrose) condition (Figure 2A, paired t-test). Similarly, the heat resistance of spores from all strains after 240 minutes in YP was statistically similar to that of dormant spores (Figure 2b), suggesting that the absence of dextrose alone causes a defect of spore activation even in presence of all other nutrients. Dextrose is therefore necessary but not sufficient for spore activation.

**Figure 2 -.**
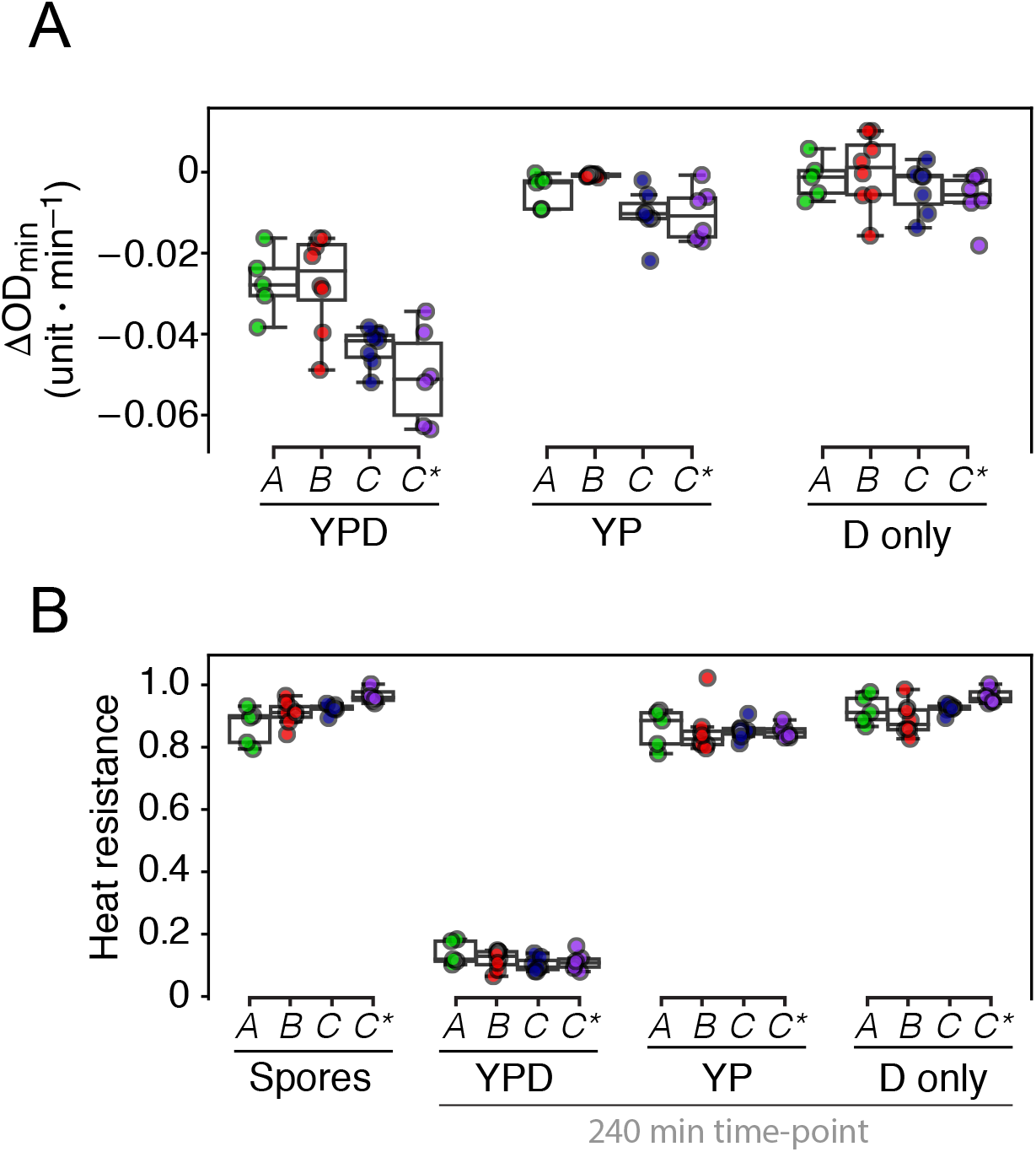
Dextrose alone is not sufficient to induce spore activation. A) Steepest decrease of OD_595_ (in units per minute) measured for each 28 strains after induction of purified spores in complete YPD, medium lacking dextrose (YP), or a 2% dextrose solution (G only). B) Heat resistance was measured for purified spores at initial time-point (Spores), or 240 minutes after induction of germination either in complete YPD, lacking dextrose medium (YP) or 2% dextrose solution (D only).

In addition, 2% dextrose solution in water (D only) was not sufficient to induce spore activation of wild *S. paradoxus* lineages. Both ΔOD_min_ and heat resistance at 240-minutes time-point values of spores in the 2 % dextrose solution were statistically similar to their respective values in YP (lacking dextrose). This result contrasts with previous reports that found dextrose to be sufficient to trigger early events in germination of *S. cerevisiæ* spores (Herman and Rine 1997). Our results show that exit from dormancy in wild *S. paradoxus* spores is not solely a response to dextrose, but requires various nutritional cues.

Dissection of a complexe medium such as YPD would be a difficult task in search of precise nutritional requirements for germination. Therefore, we tested spore germination in minimal synthetic medium (SD), whose composition can be modified in order to identify precise compounds involved in activation. Higher ΔOD_min_ was detected for purified *SpA* and *SpB* spores in minimal synthetic medium compared to YPD (Figure 3A, p=0.002 and 0.0005 respectively, Krustal-Wallis test). Moreover, heat resistance at the 240-minute time-point of *SpA* and *SpB* spores in minimal medium was comparable to resting spore values, suggesting these spores failed to activate, while *SpC* and *SpC\** spores did activate in minimal medium (Figure 3A). This is a notable difference since vegetative cells of all lineages are known to grow in minimal synthetic medium (Leducq et al 2017). It seems that compounds present in YPD but absent in minimal synthetic medium are essential cues to trigger germination of *SpA* and *SpB* spores. Extensive dissection of YPD medium could reveal the precise requirements for *SpA* and *SpB* spore activation. We could even expect such analysis to unveil further difference between *SpA* and *SpB* germination cues.

**Figure 3 -.**
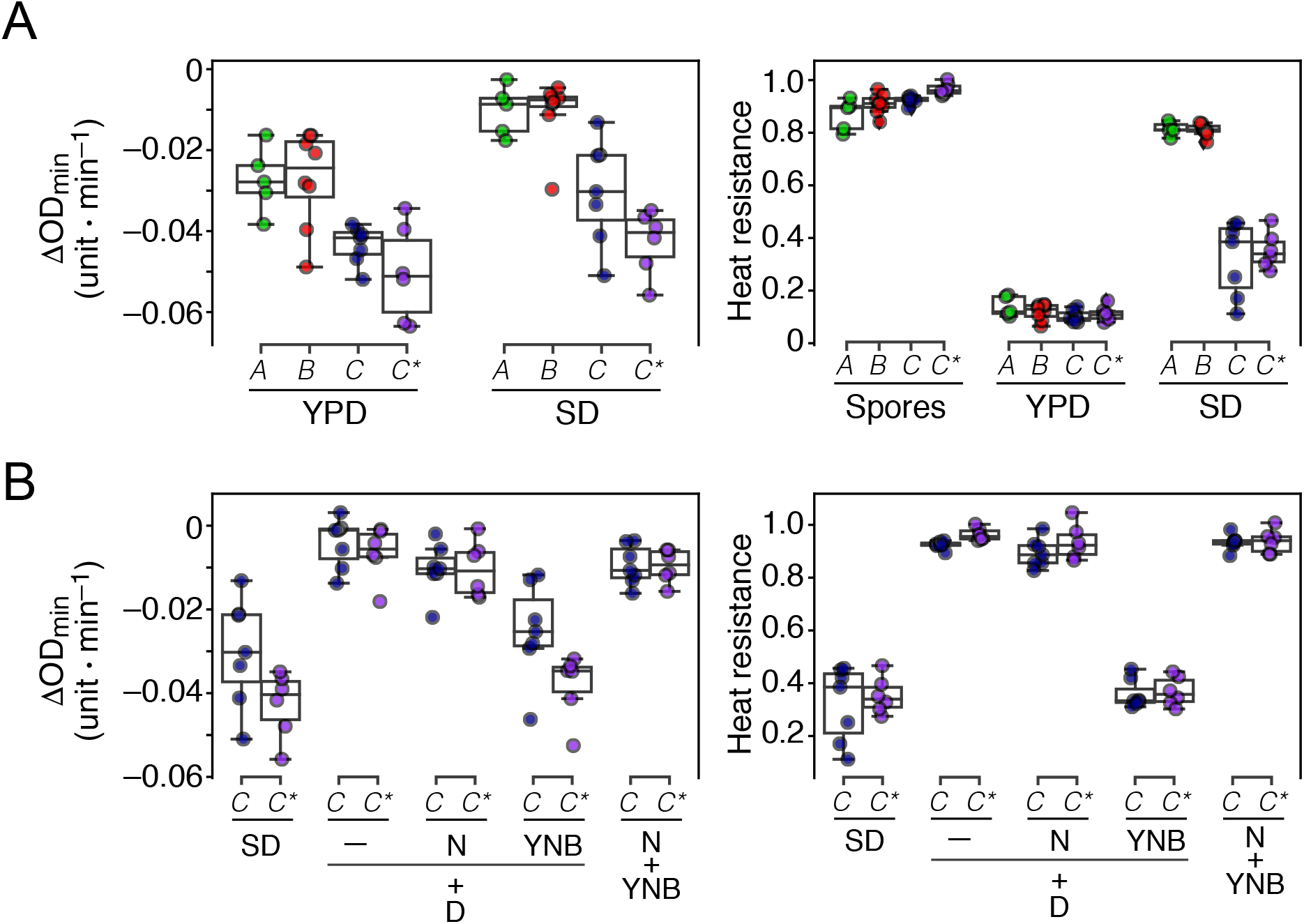
Spore activation in defined media. A) Germination of purified spores from all lineages were induced in YPD and minimal synthetic medium. Left, steepest decreases in OD_595_ (in units per minute) was measured. Right, heat resistance was measured at initial time-point (Spores), or 240 minutes after induction of germination in either YPD or minimal synthetic medium. B) *SpC* and *SpC*\* spores were inoculated in complete minimal synthetic medium, 2% dextrose without (-) or with either ammonium sulfate (N) or yeast nitrogen base (YNB), or ammonium sulfate supplemented with YNB (N+YNB). Left, steepest decrease of OD_595_ (in units per minute) measured. Right, heat resistance was measured at 240 minutes after inoculation.

Although *SpA* and *SpB* spores failed to germinate in minimal synthetic medium, *SpC* and *SpC\** lineages were valued models for further dissection of the activation signal provided by this medium. Minimal synthetic medium is composed of dextrose (D), a nitrogen source (N, ammonium sulfate) and trace elements in yeast nitrogen base (YNB, without ammonium sulfate, without amino acids). We tested the response of spores in different combinations of these components. Only the combination of dextrose and YNB produced spore activation similar to whole minimal synthetic medium, suggesting that a nitrogen source is dispensable for the exit of dormancy. In these conditions, however, spores failed to outgrow and did not complete germination to budding. Later stages of germination in *S. cerevisiæ* was previously found dependent on various nutritional conditions, including nitrogen availability (Joseph-Stauss 2007). Since we are mainly interested in the early events of germination, failure in outgrowth is not a concern.

Combination of dextrose and YNB did provide the adequate conditions to support activation of *SpC* and *SpC*\* spores. YNB is composed of various compounds that support fungal growth, which include vitamins, growth factors, and inorganic salts, providing essential metal ions (Figure 4A). We tested whether the vitamins (Organic, Figure 4) or inorganic salts is the source for spore activation signals. *SpC* and *SpC*\* spores in 2% dextrose with all the inorganic salts did activate as well as with complete YNB, while the vitamins and growth factor alone with dextrose failed to activate. This result narrows candidate compounds down to 11 inorganic salts to test for their ability to induce germination. Potassium phosphate (KH_2_PO_4_) alone with 2% dextrose induced spore activation as well as complete YNB, while the addition of the other ten compounds did not induce a strong activation. Addition of magnesium sulfate (MgSO_4_) did lower significantly heat resistance (paired t-test, p < 0.05) and induced OD_595_ decrease. However, only the omission of potassium phosphate in YNB abolished its ability to induce spore activation in addition to dextrose (Figure S3). This result is indicative of the importance of potassium phosphate to exit dormancy.

**Figure 4 -.**
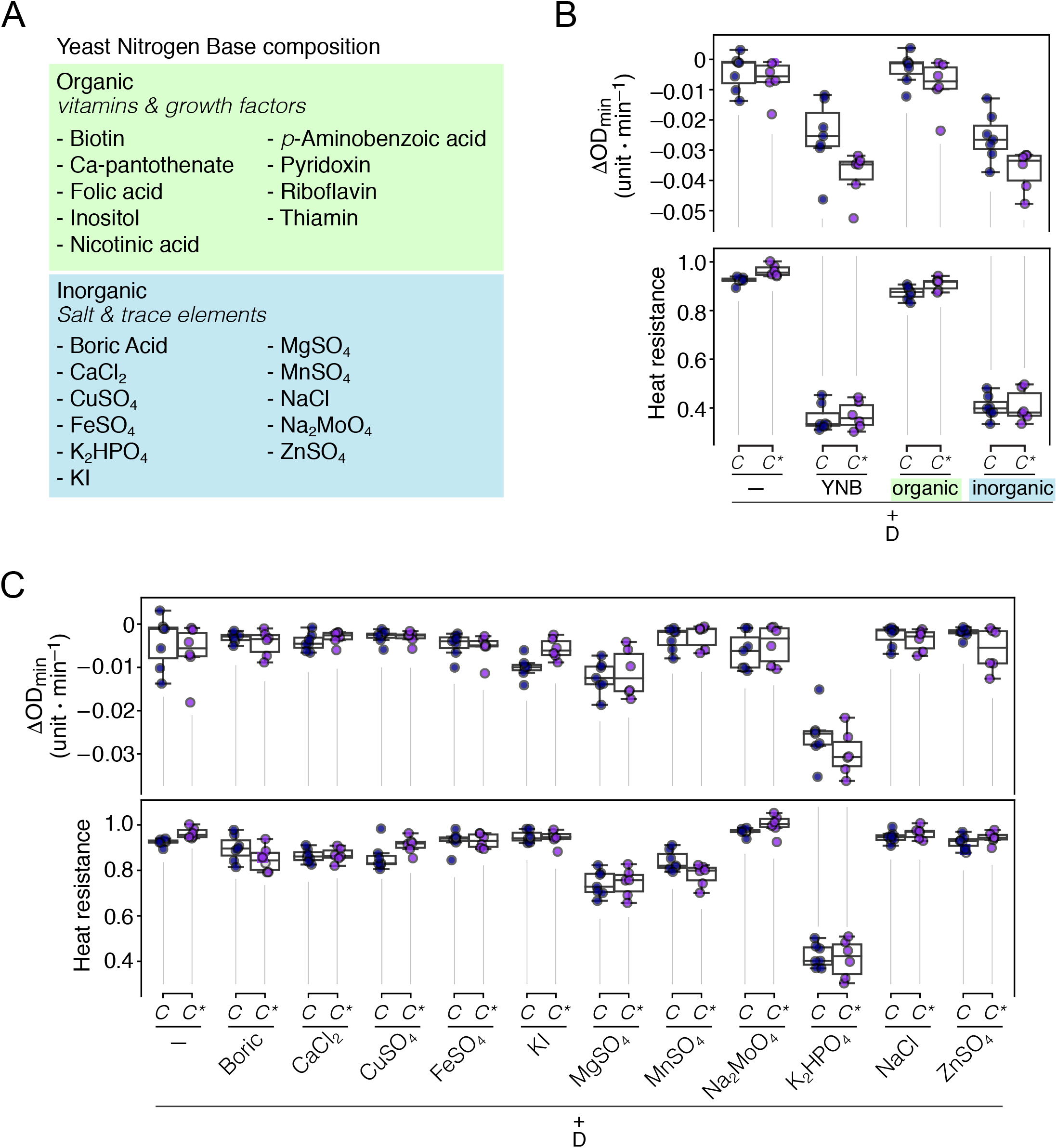
Potassium phosphate triggers spore activation in the presence of dextrose in *SpC* and *SpC*\*. A) Yeast nitrogen base (YNB) components were divided in either organic (including vitamins and growth factors) or inorganic parts (including salts and trace metal ions). B) *SpC* and *SpC*\* spores were inoculated in 2% dextrose without (-) or with either complete YNB, organic, or inorganic compounds of YNB. Top, steepest decrease of OD_595_ (in units per minute) measured. Bottom, heat resistance was measured at 240 minutes after inoculation. C) *SpC* and *SpC\** spores were inoculated in 2% dextrose without (-) or with each inorganic component. Top, steepest decrease of OD_595_ (in units per minute) measured. Bottom, heat resistance was measured at 240 minutes after inoculation.

While complete YNB with dextrose has a pH of 4.8, omission of KH_2_PO_4_ increases pH to 6.8. Therefore, we tested whether it is the phosphate itself or the buffered conditions that drive spore activation. Substitution of KH_2_PO_4_ with K_2_HPO_4_ in YNB results in a medium with pH 8.5, higher than original YNB; yet, spores activated as well as when using original YNB. Additionally, substitution of potassium phosphate with acetate buffer (NaOAc, Figure S3), with a pH of 4.8, was not a proper condition for spore activation. Together, these results show that potassium phosphate is an essential compound for *SpC* and *SpC*\* activation.

## Conclusion

Our findings highlight that dextrose is essential, but not sufficient to trigger spore activation in *S. paradoxus*, meaning that additional nutrients must contribute to signal germination. Dissection of germination medium allowed the identification of inorganic phosphate as an additional environmental cue that signal germination induction in *SpC* and *SpC*\* lineages. The contribution of phosphate to spore germination has also been characterized in the yeast *Pichia pastoris*. The presence of both dextrose and phosphate in the environment of these spores is required for maximal trehalase activity (Thevelein et al., 1982). Because trehalase catalyzes the breakdown of trehalose, an essential step of spore germination (Thevelein et al., 1984), this observation suggests a potential mechanism of action for phosphate. Although the signal of dextrose and phosphate seems specific for *SpC* and *SpC*\* spore activation among *S. paradoxus* lineages in North America, it may actually be shared by other fungal species. However, the exact mechanism of action requires to be examined.

The four lineages of *S. paradoxus* studied here divergerged relatively recently, with *SpC* and *SpC*\* being the most closely related species, with most divergence due to *SpC*\* having ~5% of its genome introgressed from *SpB* (*Leducq et al*., 2016). The fact that both *SpC* and *SpC*\* respond to similar signals (inorganic phosphate) but that are different from those to which *SpA* and *SpB* respond suggest that it was recently acquired after the split with *SpB*. The distinct spore activation signals between *SpB* and *SpC/SpC*\* provides a potential ecologically based mechanisms for the reproductive isolation of these lineages. The adaptive significance of responding to phosphate and dextrose for these lineages remains to be investigated. One possibility is that phosphate is particularly limiting in its environment and spore activation needs to take place only if critical concentrations are met. The key elements that trigger spore activation for *SpA* and *SpB*, which is present in YPD but absent in synthetic medium, remains to be identified and they may not be the same for both lineages. Further work on this aspect of yeast ecology remains to be done and the tools and approaches we have developed here will help such future investigations.

## Acknowledgements

We thank Alexandre K Dubé and Philippe Després for their comments on the manuscript, and Alexandre K Dubé and Isabelle Gagnon-Arsenault for their support in the laboratory. This work was supported by a NSERC Discovery Grant to CRL. CRL holds the Canada Research Chair in Cellular Synthetic and Systems Biology.

## Supplementary figure legends

**Figure S1 -.**
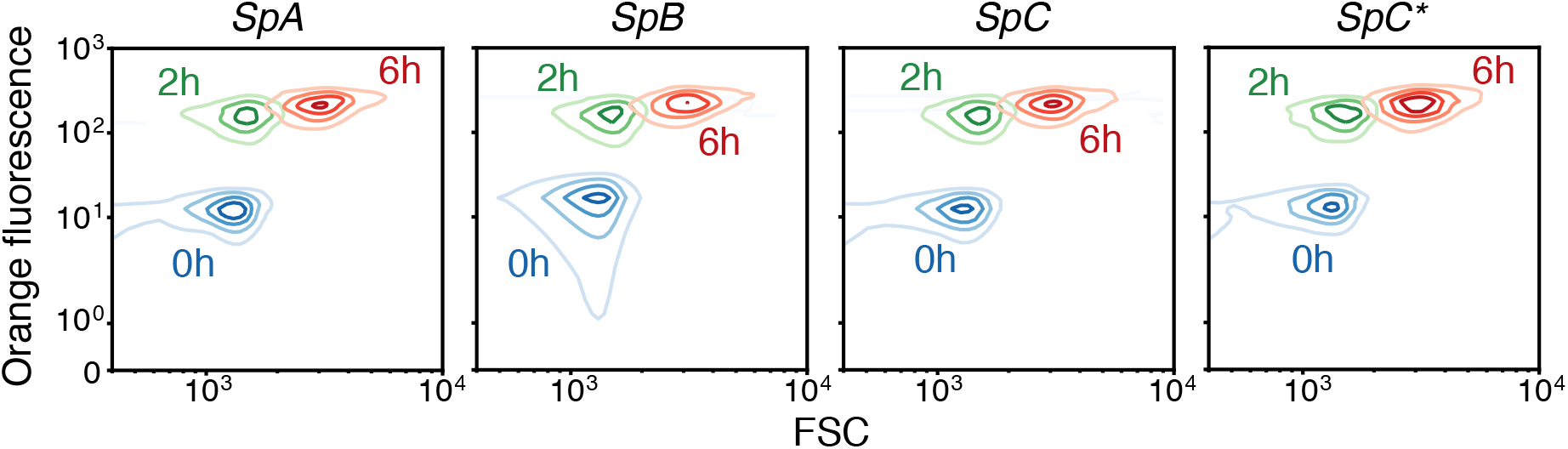
Flow cytometry analysis of spore germination through time. Germination of spores from *SpA* (LL12_001), *SpB* (LL12_009), *SpC* (LL11_011) and *SpC*\* (LL11_006) strains were induced in YPD and analysed by flow cytometry. At initial time-point (0h, blue), 2h (green) and 6h (red) after induction cells were analyzed by flow cytometry. Forward scatter (FCS) and orange fluorescent signals are shown. Signals for 5000 events per time-points are plotted here as Kernel density plot. Each of the four estimated density levels shown represents 25% of events.

**Figure S2 -.**
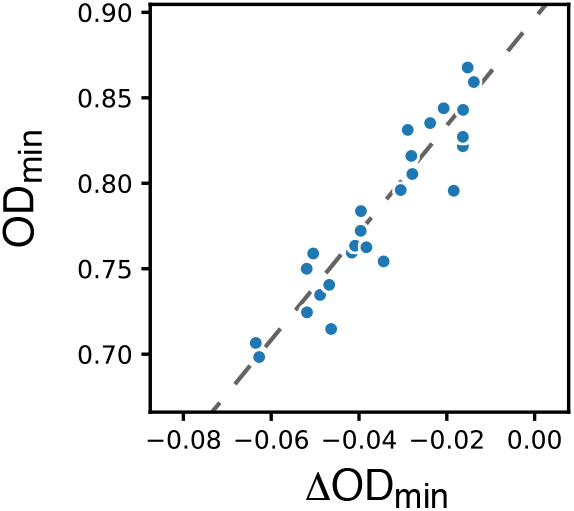
Correlation between steepest decrease in OD595 and minimal OD595 reached. Germination of spores from 28 strains was induced in YPD, and the steepest decrease value in OD_595_ and minimal OD_595_ value reached for each strain are plotted here. Shown are representative data of 4 replicates.

**Figure S3 -.**
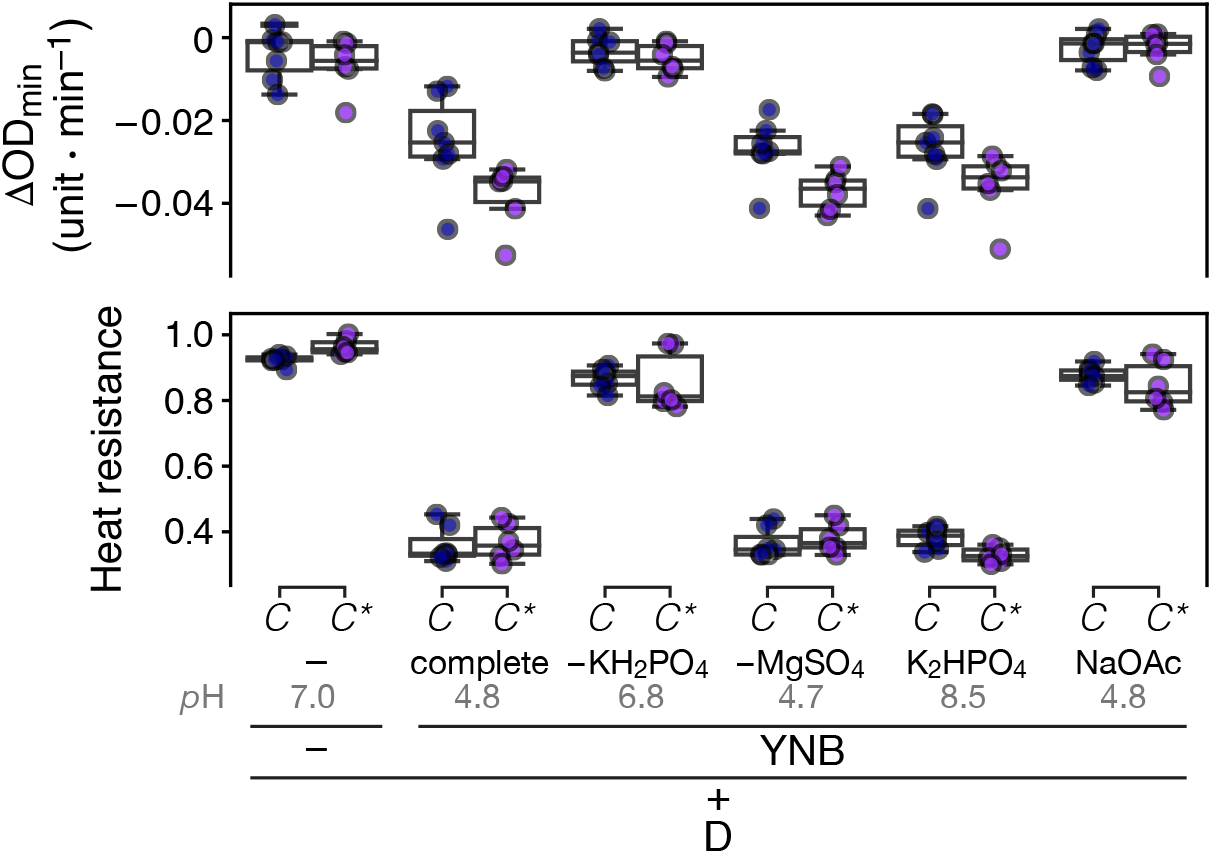
Spore activation is triggered by phosphate compounds and not by pH conditions. *SpC* and *SpC\** spores were inoculated in 2% dextrose solution without (-) or with either complete YNB, YNB lacking potassium phosphate, YNB lacking magnesium sulfate, YNB in which potassium phosphate monobasic was substituted with potassium phosphate dibasic, or YNB in which potassium phosphate monobasic was substituted with acetate buffer (pH 4.8). Top, steepest decrease of OD_595_ (in units per minute) measured. Bottom, heat resistance was measured at 240 minutes after inoculation. pH of each media measured at 25°C are shown.

## References

Allen, P. J. (1965). Metabolic Aspects of Spore Germination in Fungi. Annual Review of Phytopathology, 3(1), 313–342.

Bagga, S., & Straney, D. (2000). Modulation of cAMP and phosphodiesterase activity by flavonoids which induce spore germination of Nectria haematococca MP VI (Fusarium solani). Physiological and Molecular Plant Pathology, 56(2), 51–61.

Botts, M. R., & Hull, C. M. (2010). Dueling in the lung: how Cryptococcus spores race the host for survival. Current Opinion in Microbiology, 13(4), 437–442.

Calhim, S., Halme, P., Petersen, J. H., Læssøe, T., Bässler, C., & Heilmann-Clausen, J. (2018). Fungal spore diversity reflects substrate-specific deposition challenges. Scientific Reports, 8(1), 53–56.

Charron, G., Leducq, J.-B., Bertin, C., Dubé, A. K., & Landry, C. R. (2014). Exploring the northern limit of the distribution of Saccharomyces cerevisiae and Saccharomyces paradoxus in North America. FEMS Yeast Research, 14(2), 281–288.

Charron, G., Leducq, J.-B., & Landry, C. R. (2014). Chromosomal variation segregates within incipient species and correlates with reproductive isolation. Molecular Ecology, 23(17), 4362–4372.

Coluccio, A. E., Rodriguez, R. K., Kernan, M. J., & Neiman, A. M. (2008). The yeast spore wall enables spores to survive passage through the digestive tract of Drosophila. PloS One, 3(8), e2873.

d’Enfert, C. (1997). Fungal Spore Germination: Insights from the Molecular Genetics of Aspergillus nidulans and Neurospora crassa. Fungal Genetics and Biology: FG & B, 21(2), 163–172.

Eberlein, C., Hénault, M., Fijarczyk, A., Charron, G., Bouvier, M., Kohn, L. M., Anderson, J. B., & Landry, C. R. (2019). Hybridization is a recurrent evolutionary stimulus in wild yeast speciation. Nature Communications, 10(1), 923.

French, R. C. (1992). Volatile Chemical Germination Stimulators of Rust and Other Fungal Spores. Mycologia, 84(3), 277–288.

Fröhlich-Nowoisky, J., Pickersgill, D. A., Després, V. R., & Pöschl, U. (2009). High diversity of fungi in air particulate matter. Proceedings of the National Academy of Sciences of the United States of America, 106(31), 12814–12819.

Greig, D. (2009). Reproductive isolation in Saccharomyces. Heredity, 102(1), 39–44.

Halbwachs, H., BÃssler C., & Brandl, R. (2015). Spore wall traits of ectomycorrhizal and saprotrophic agarics may mirror their distinct lifestyles. Fungal Ecology, 17. https://doi.org/10.1016/j.funeco.2014.10.003

Hatanaka, M., & Shimoda, C. (2001). The cyclic AMP/PKA signal pathway is required for initiation of spore germination in Schizosaccharomyces pombe. Yeast, 18(3), 207–217.

Hayer K., Stratford, M., & Archer, D. B. (2013). Structural features of sugars that trigger or support conidial germination in the filamentous fungus Aspergillus niger. Applied and Environmental Microbiology, 79(22), 6924–6931.

Hénault, M., Eberlein, C., Charron, G., Durand, É., Nielly-Thibault, L., Martin, H., & Landry, C. R. (2017). Yeast Population Genomics Goes Wild: The Case of Saccharomyces paradoxus. In Population Genomics: Microorganisms (pp. 207–230). Springer.

Herman, P. K., & Rine, J. (1997). Yeast spore germination: a requirement for Ras protein activity during re-entry into the cell cycle. The EMBO Journal, 16(20), 6171–6181.

Huang, M., & Hull, C. M. (2017). Sporulation: how to survive on planet Earth (and beyond). Current Genetics, 63(5), 831–838.

Joseph-Strauss, D., Zenvirth, D., Simchen, G., & Barkai, N. (2007). Spore germination in Saccharomyces cerevisiae: global gene expression patterns and cell cycle landmarks. Genome Biology, 8(11), R241.

Kloimwieder, A., & Winston, F. (2011). A Screen for Germination Mutants in Saccharomyces cerevisiae. G3, 1(2), 143–149.

Kono, K., Matsunaga, R., Hirata, A., Suzuki, G., Abe, M., & Ohya, Y. (2005). Involvement of actin and polarisome in morphological change during spore germination of Saccharomyces cerevisiae. Yeast, 22(2), 129–139.

Leducq, J.-B., Nielly-Thibault, L., Charron, G., Eberlein C., Verta, J.-P., Samani, P., Sylvester, K., Hittinger, C. T., Bell, G., & Landry, C. R. (2016). Speciation driven by hybridization and chromosomal plasticity in a wild yeast. Nature Microbiology, 1, 15003.

Maclean, C. J., & Greig, D. (2008). Prezygotic reproductive isolation between Saccharomyces cerevisiae and Saccharomyces paradoxus. BMC Evolutionary Biology, 8, 1.

Murphy, H. A., & Zeyl, C. W. (2012). PREZYGOTIC ISOLATION BETWEEN SACCHAROMYCES CEREVISIAE AND SACCHAROMYCES PARADOXUS THROUGH DIFFERENCES IN MATING SPEED AND GERMINATION TIMING. In Evolution (Vol. 66, Issue 4, pp. 1196–1209). https://doi.org/10.1111/j.1558-5646.2011.01516.x

Neiman, A. M. (2005). Ascospore formation in the yeast Saccharomyces cerevisiae. Microbiology and Molecular Biology Reviews: MMBR, 69(4), 565–584.

Neiman, A. M. (2011). Sporulation in the budding yeast Saccharomyces cerevisiae. Genetics, 189(3), 737–765.

Osherov, N. (2009). Conidial Germination in Aspergillus fumigatus. In Aspergillus fumigatus and Aspergillosis (pp. 131–142). asm Pub2Web.

Plante, S., & Labbé, S. (2019). Spore Germination Requires Ferrichrome Biosynthesis and the Siderophore Transporter Str1 in Schizosaccharomyces pombe. Genetics, 211(3), 893–911.

Podila, G. K., Rogers, L. M., & Kolattukudy, P. E. (1993). Chemical Signals from Avocado Surface Wax Trigger Germination and Appressorium Formation in Colletotrichum gloeosporioides. Plant Physiology, 103(1), 267–272.

Pringle, A., Vellinga, E., & Peay, K. (2015). The shape of fungal ecology: does spore morphology give clues to a species’ niche? Fungal Ecology, 100(17), 213–216.

Rieux, A., Soubeyrand, S., Bonnot, F., Klein, E. K., Ngando, J. E., Mehl, A., Ravigne, V., Carlier, J., & de Lapeyre de Bellaire, L. (2014). Long-distance wind-dispersal of spores in a fungal plant pathogen: estimation of anisotropic dispersal kernels from an extensive field experiment. PloS One, 9(8), e103225.

Rousseau, P., Halvorson, H. O., Bulla, L. A., Jr, & St Julian, G. (1972). Germination and outgrowth of single spores of Saccharomyces cerevisiae viewed by scanning electron and phase-contrast microscopy. Journal of Bacteriology, 109(3), 1232–1238.

Sando, N., Oguchi, T., Nagano, M., & Osumi, M. (1980). Morphological changes in ascospores of Saccharomyces cerevisiae during aerobic and anaerobic germination. The Journal of General and Applied Microbiology, 26(6), 403–412.

Schmit, J. C., & Brody, S. (1975). Neurospora crassa conidial germination: role of endogenous amino acid pools. Journal of Bacteriology, 124(1), 232–242.

Shimoda, C. (1980). Differential effect of glucose and fructose on spore germination in the fission yeast, Schizosaccharomyces pombe. Canadian Journal of Microbiology, 26(7), 741–745.

Smits, G. J., van den Ende, H., & Klis, F. M. (2001). Differential regulation of cell wall biogenesis during growth and development in yeast. Microbiology, 147(Pt4), 781–794.

Sprouffske, K., & Wagner, A. (2016). Growthcurver: an R package for obtaining interpretable metrics from microbial growth curves. BMC Bioinformatics, 17, 172.

Stelkens, R. B., Miller, E. L., & Greig, D. (2016). Asynchronous spore germination in isogenic natural isolates of Saccharomyces paradoxus. FEMS Yeast Research, 16(3). https://doi.org/10.1093/femsyr/fow012

Thevelein, J. M., den Hollander, J. A., & Shulman, R. G. (1982). Changes in the activity and properties of trehalase during early germination of yeast ascospores: Correlation with trehalose breakdown as studied by in vivo13C NMR. Proceedings of the National Academy of Sciences of the United States of America, 79(11), 3503–3507.

Thevelein, J. M., den Hollander, J. A., & Shulman, R. G. (1984). Trehalase and the control of dormancy and induction of germination in fungal spores. Trends in Biochemical Sciences, 9(11), 495–497.

Tong, A. H. Y., & Boone, C. (2006). Synthetic genetic array analysis in Saccharomyces cerevisiae. Methods in Molecular Biology, 313, 171–192.

Tsai, I. J., Bensasson, D., Burt, A., & Koufopanou, V. (2008). Population genomics of the wild yeast Saccharomyces paradoxus: Quantifying the life cycle. Proceedings of the National Academy of Sciences of the United States of America, 105(12), 4957–4962.

Van Laere, A., Van Schaftingen, E., & Hers, H. G. (1983). Fructose 2,6-bisphosphate and germination of fungal spores. Proceedings of the National Academy of Sciences of the United States of America, 80(21), 6601–6605.

van Leeuwen, M. R., Krijgsheld, P., Bleichrodt, R., Menke, H., Stam, H., Stark, J., Wösten, H. A. B., & Dijksterhuis, J. (2013). Germination of conidia of Aspergillus niger is accompanied by major changes in RNA profiles. Studies in Mycology, 74(1), 59–70.

Walsh, N. M., Botts, M. R., McDermott, A. J., Ortiz, S. C., Wüthrich, M., Klein, B., & Hull, C. M. (2019). Infectious particle identity determines dissemination and disease outcome for the inhaled human fungal pathogen Cryptococcus. PLoS Pathogens, 15(6), e1007777.

